# MIXTURE: an improved algorithm for immune tumor microenvironment estimation based on gene expression data

**DOI:** 10.1101/726562

**Authors:** Elmer A. Fernández, Yamil D. Mahmoud, Florencia Veigas, Darío Rocha, Mónica Balzarini, Hugo D. Lujan, Gabriel A. Rabinovich, M. Romina Girotti

## Abstract

RNA sequencing has proved to be an efficient high-throughput technique to robustly characterize the presence and quantity of RNA in tumor biopsies at a given time. Importantly, it can be used to computationally estimate the composition of the tumor immune infiltrate and to infer the immunological phenotypes of those cells. Given the significant impact of anti-cancer immunotherapies and the role of the associated immune tumor microenvironment (ITME) on its prognosis and therapy response, the estimation of the immune cell-type content in the tumor is crucial for designing effective strategies to understand and treat cancer. Current digital estimation of the ITME cell mixture content can be performed using different analytical tools. However, current methods tend to over-estimate the number of cell-types present in the sample, thus under-estimating true proportions, biasing the results. We developed MIXTURE, a noise-constrained recursive feature selection for support vector regression that overcomes such limitations. MIXTURE deconvolutes cell-type proportions of bulk tumor samples for both RNA microarray or RNA-Seq platforms from a leukocyte validated gene signature. We evaluated MIXTURE over simulated and benchmark data sets. It overcomes competitive methods in terms of accuracy on the true number of present cell-types and proportions estimates with increased robustness to estimation bias. It also shows superior robustness to collinearity problems. Finally, we investigated the human immune microenvironment of breast cancer, head and neck squamous cell carcinoma, and melanoma biopsies before and after anti-PD-1 immunotherapy treatment revealing associations to response to therapy which have not seen by previous methods.

## Introduction

Recruitment and activation of immune cells to the tumor microenvironment are closely associated with clinical outcome (Mantovani et al. 2008; Gajewski et al. 2013). The malignant phenotype of cancer is defined not only by the intrinsic activities of cancer cells but also by components in the tumor microenvironment, especially tumor-infiltrating immune cells. The immune infiltrate in both primary and metastatic sites correlates closely with patient prognosis and survival in several cancer types (Fridman et al. 2017). Cytotoxic CD8 T cells are mostly linked to prolonged survival (Galon et al. 2006). For example, a high density of CD8+ T cells is predictive of a longer overall survival duration in patients with breast cancer, head and neck cancer and colorectal cancer (CRC) with hepatic and/or lung metastases (Fridman et al. 2017), but also predicts poor overall survival in patients with lung metastases from clear cell renal cell carcinoma (ccRCC) (Giraldo et al. 2015).

Immunosuppressive T-regulatory cells are associated with shorter survival in non-small cell lung cancer (NSCLC), pancreatic and breast cancer, RCC, hepatocellular carcinoma (HCC) and melanoma (Fridman et al. 2012; Fridman et al. 2017). In most tumors, macrophages mainly display an M2 phenotype, generally favoring the growth and development of an invasive and pro-angiogenic phenotype. The density of such macrophages correlates with a poor prognosis among patients with breast, bladder, ovarian, gastric, or prostate cancers, RCC and melanoma (Fridman et al. 2017). Based on the protumorigenic properties of M2-macrophages, several clinical trials are currently testing the effect of macrophage inhibitors in combination with PD-1/PD-L1- and CTLA-4-targeting immunotherapies (Yang and Zhang 2017). Conversely, a high density of M1-macrophages has been correlated with a favorable prognosis among patients with ovarian and gastric cancer, NSCLC and HCC (Fridman et al. 2017).

The development of new and effective anti-cancer immune therapies based on immune checkpoint blockade (ICB) has revolutionized the treatment of cancer due to the significant improvement of patient survival in several indications (Couzin-Frankel 2013; Callahan et al. 2016). However, since therapy remains ineffective for a substantial number of patients, there is an urgent clinical need to identify and develop predictive biomarkers of response to ICB both to foster precision immunotherapy and to understand and overcome the mechanisms of resistance. Retrospective analyses of patient immune infiltrate revealed that tumor–immune phenotypes influence response to immune checkpoint inhibitors (Havel et al. 2019). Hence, the characterization of the immune tumor microenvironment (ITME) has been suggested as a predictor of therapy response and outcome (Bense et al. 2017). Therefore, its analysis represents a possible, reliable biomarker of response to therapy (Fridman et al. 2012; Herbst et al. 2014; Tumeh et al. 2014; Mariathasan et al. 2018; Havel et al. 2019).

Several computational approaches and algorithms have made possible to systematically infer the ITME composition levels, i.e. digital cytometry, in large-scale cancer patient datasets using gene expression information (Gaujoux and Seoighe 2013; Yoshihara et al. 2013; Cesano 2015; Newman et al. 2015; Becht et al. 2016; Aran et al. 2017). Thanks to these computational approaches, the immune cell content, represented by a reference molecular signature of known cell-types gene expression profiles, can be computationally dissected from a subject gene expression profile by means of statistical and/or machine learning linear methods (Abbas et al. 2009; Newman et al. 2015; Li et al. 2017; Monaco et al. 2019; Newman et al. 2019). A recent study in pan-cancer samples has demonstrated that the presence of T-lymphocyte and B-lymphocyte signatures, estimated by a digital cytometry method, is associated with a favorable prognosis (Gentles et al. 2015). The presence of specific plasma-cell signatures was also associated with a good prognosis. Among T-cell subsets, the presence of regulatory T-cells (Tregs) indicated a poor prognosis, whereas a signature that included γδT-cells constituted the most influential factor indicating a favorable prognosis. Signatures including myeloid cells (macrophages, neutrophils, eosinophils granulocytes and dendritic cells), NK cells, and also those including memory B cells were all associated with a poor prognosis (Gentles et al. 2015). Although these works show the potential of deconvolution methods to infer the immune infiltrate, current methods have been compared with highly controversial results regarding their efficacy (Li et al. 2017; Hunt et al. 2018). Here it is shown that current competitive state of the art methods (Abbas et al. 2009; Newman et al. 2015; Hunt et al. 2018; Monaco et al. 2019) tend to overestimate the number of present cell types, providing inaccurate and biased proportion immune estimates as well as lack of robustness to multicollinearity of the cell-types profiles present in the signature matrix and prediction dependent bias. These problems may negatively impact their reliability as therapy response biomarkers, as well as their use to infer cell-type gene expression profiles from bulk sample cohorts (Newman et al. 2019).

To overcome such limitations, we developed a new digital cytometry method, MIXTURE, utilizing a v-SVR-based noise constrained Recursive Feature Selection algorithm. We compared MIXTURE against the current state of the art methods (Abbas et al. 2009; Newman et al. 2015; Hunt et al. 2018; Monaco et al. 2019) to estimate 22 mature human hematopoietic populations and activation states cell-type proportions [LM22 molecular signature from (Newman et al. 2015)] over simulated and flow cytometry derived cell-type proportions data. We have also analyzed patients’ data from breast cancer (BRCA - TCGA), head and neck squamous cell carcinoma (HNSCC) with known survival outcome, and melanoma patient biopsies with known response to immunotherapy. Our analyses show that MIXTURE outperforms the competing methods and helps to estimate the immune cell landscape associated with outcome and response to immune checkpoint blockade accurately.

## Results

### Algorithm Overview

The linear deconvolution of the cell types present in a gene signature matrix (**X**), holding *N* genes for *k* cell types, associated with the components of a mixture of cell types present in a tumor gene expression profile (***Y***), involves solving the following regression model equation **Y=X ⋅ B^T^**, where the proportions for all cell types in the mixture sample is represented by the column tor **B** = {β_*j*_≥ 0 ^ ∑ β_*j*_ = 1 ∀*j* = 1,…,*k*} (Avila Cobos et al. 2018). Our newly developed algorithm, MIXTURE, provides accurate estimates of the number of cell types present in the samples with a consequent improvement in the estimated proportion cell levels. MIXTURE define falsely detected cell types based on a floating-point error constrain in addition to a recursive feature selection process to iteratively remove such false detected cell types upon a v-SVR approach as a regression function similar to the one used by CIBERSORT (see supplementary material). Source code of MIXTURE, as well as all the described evaluations, are available at https://github.com/elmerfer/MIXTURE.App

### Simulated scenarios

We compared MIXTURE performance against state of the art algorithms (i.e ABBAS, ABIS, CIBERSORT and DTANGLE) over simulated, flow cytometry derived benchmark data and cancer data sets as follows:

#### Scenario 1

the algorithms were tested using ***Y*** as: S1.1) the single pure cell types present in the LM22 signature matrix (i.e. the same cell types of the immune cell-type molecular signature ***X*** were alternatively used as the observed expression profile; i.e., ***Y***=***X***_k_, ***X***_k_ being a column of the gene signature matrix LM22 from (Newman et al. 2015), S1.2) Simulated mixture of cell types without noise: between 2 and 8 cell types of the immune cell-type molecular signature LM22 were randomly selected to simulate ***Y***, ***X*** being the LM22 matrix. At each simulation run, the cell type proportions were sampled from a uniform [0.2, 1] distribution and normalized to satisfy the sum-to-one constrain. Then the cell-types were multiplied by the sampled proportions and summed to build the simulated ***Y***. The process was simulated 1000 times. For the Scenario S1.3, the simulated mixture of cell-types with noise was same as S1.2 but adding a noise vector to ***Y***; this noise vector is composed of a random sample of N (i.e., the number of genes in the signature matrix) values of gene expressions drawn from LM22.

#### Scenario 2

S2.1) The same data from Newman et al., 2015 and Hunt et al, 2018 were used. The Newman Follicular Lymphoma (FL) data set, generated by taking lymph node biopsy samples and enumerating the immune cell sub-types using flow cytometry (Newman et al. 2015). In this case, the process identified 3 leukocyte types (B, CD8, and CD4) in various proportions across 14 patient samples. Since LM22 contains 22 cell-types, the proportions of those leukocytes types were summarized according to (Newman et al. 2015). S2.2) The Newman Peripheral Blood Mononuclear Cells (PBMC) data, generated from blood samples from 22 adults where the proportion of nine leukocytes were determined by flow cytometry.

#### Scenario 3

The algorithms were applied over real data to evaluate: S3.1) the 1095 Breast Cancer TCGA biopsies (Cancer Genome Atlas 2012), downloaded from TCGA repository site in November 2018 (see supplementary material). These RNA-Seq data used to feed the PBCMC R library (Fresno et al. 2017). In order to confidently assign subjects to the PAM50 intrinsic breast cancer subtypes S3.2) Transcriptomic samples of 81 head and neck squamous cell carcinoma (HNSCC) primary tumors (Rickman et al. 2008) used to investigate the propensity for subsequent distant metastasis from patients initially treated by surgery that developed or not-developed metastases as the first recurrent event. S3.3) Transcriptomic tumor samples from 68 patients with advanced melanoma, who progressed on the anti-CTLA-4 drug ipilimumab or were ipilimumab naive, before and after they commenced anti-PD-1 therapy with nivolumab (Riaz et al. 2017).

### Comparing methods over simulated data

When cell-type proportions were estimated under scenario S1.1, the expected result was the identification of a single cell type in the sample and the remaining ones should be determined as null. The CIBERSORT tool estimated between one (23% of the cases) and 8 cell-types present in the sample. This tool is affected by floating-point errors since those regression coefficients associated with false cell-type detection ranged between 1e^−4^ and 2e^−4^. The ABBAS method estimated between 6 and 9 cell-types. The range of those falsely detected cell type coefficients was between 2e-5 and 0.32, also suggesting sensitivity to floating-point errors. The DTANGLE method always estimated 22 cell-types with a range for falsely detected cell-type coefficients between 5e^−4^ and 0.35. The ABIS method does not constraint 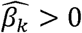 (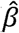 *estimated coefficient*), so those 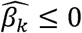 were considered as null cell-types. In this sense, it estimated between 10 to 13 cell-types and the falsely detected proportions range between 2e^−5^ to 0.25 for those estimated 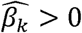 and between −0.21 to 0.25 for all estimated 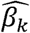 (i.e. truly and falsely detected cell types). All these methods overestimated the number of cell types in a given sample; consequently, they underestimated the true proportion of the real cell type, except the ABIS, which may over and/or underestimate, since it does not satisfies the non-negativity and sum-to-one constraints (Avila Cobos et al. 2018). Also, high regression coefficients associated with false cell-type detection by ABBAS, ABIS and DTANGLE mimic correlation coefficients patterns of the molecular signature matrix (see Supplementary Figure 1), suggesting collinearity problems affecting their estimations. On the contrary, MIXTURE indicated the presence of only one cell type in all the simulated samples.

In Figure 1 it is represented the estimated number of cell types, for the simulated cell-type mixture samples, in both scenarios S1.2 and S1.3 (without noise – left panel and with noise – right panel). It can be observed that ABBAS, ABIS and DTANGLE methods overestimate the number of cell-types; in particular, DTANGLE always provides 22 cell-types. The CIBERSORT method provides accurate estimated coefficients for mixtures samples without noise (S1.2), overestimating mainly for mixture samples composed with less than four cell-types. However, it significantly over-estimated the number of cell-types for mixture samples with noise compared against MIXTURE (Wilcoxon paired test, p < 0.001) which provides much more accurate estimates in any simulated scenario, thus resulting the method less affected by noise.

**Figure 1.**
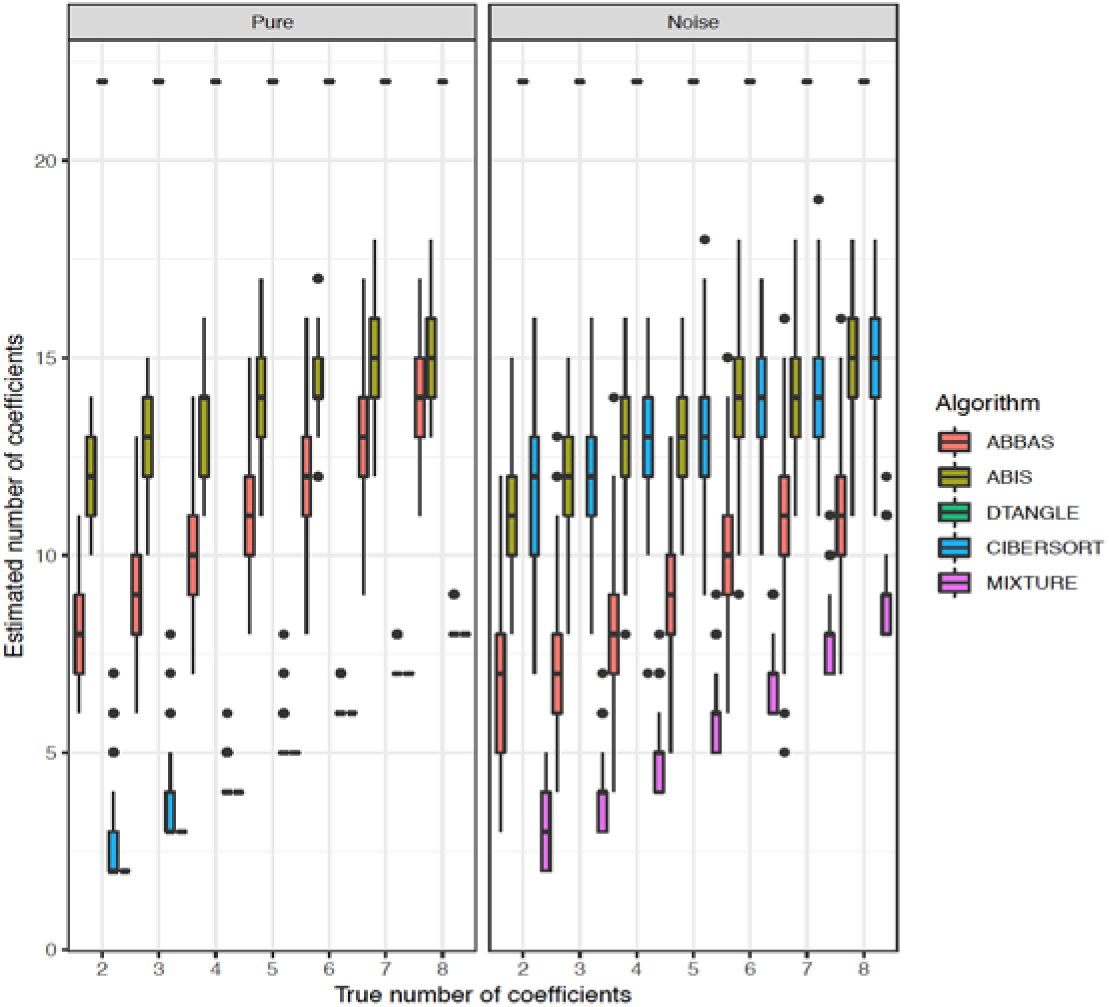
Distribution of the estimated number of coefficients (proportions) for simulated mixtures without added noise (Pure) and with it (left and right panels, respectively). It can be observed that DTANGLE consistently estimates 22 cell-types independently of the number of simulated cell-types (no boxplot is shown but overlapped points at “Estimated number of coefficients” = 22). MIXTURE provides the true number of cell-types when feeding with pure samples (without noise). CIBERSORT performs similar to MIXTURE for pure samples with more than three mixed cell-types. For noisy mixtures, MIXTURE always provides more accurate estimates of the true number of cell-types.

In Figure 2A, Bland-Altman plots represent the difference between estimated and simulated (true) coefficients. All the evaluated methods tend to underestimate the true simulated proportions (−0.039 ± 0.059, −0.021 ± 0.048, −0.113±0.113, −0.011±0.014, −0.002±0.007) for ABBAS, ABIS, DTANGLE, CIBERSORT and MIXTURE respectively. However, MIXTURE shows a lesser bias with the lesser standard deviation of the errors. Also, it can be observed that all the methods except MIXTURE show increased underestimation bias towards high proportion values. When evaluating the simulated null coefficients (i.e. cell-types not present in the sample are represented by), all methods overestimate extra proportions related to false cell-types identification. The DTANGLE method shows the highest false proportion estimates (maximum value of 0.94), followed by ABBAS and ABIS (0.48), CIBERSORT (0.15) and MIXTURE (0.11). Thus, both v-SVR based methods outperform the rest, but the noise constrained RFE algorithm implemented in MIXTURE significantly improves the accuracy of the estimated coefficients compared to CIBERSORT (Wilcoxon paired test p < 0.001, See Supplementary Material for details).

**Figure 2.**
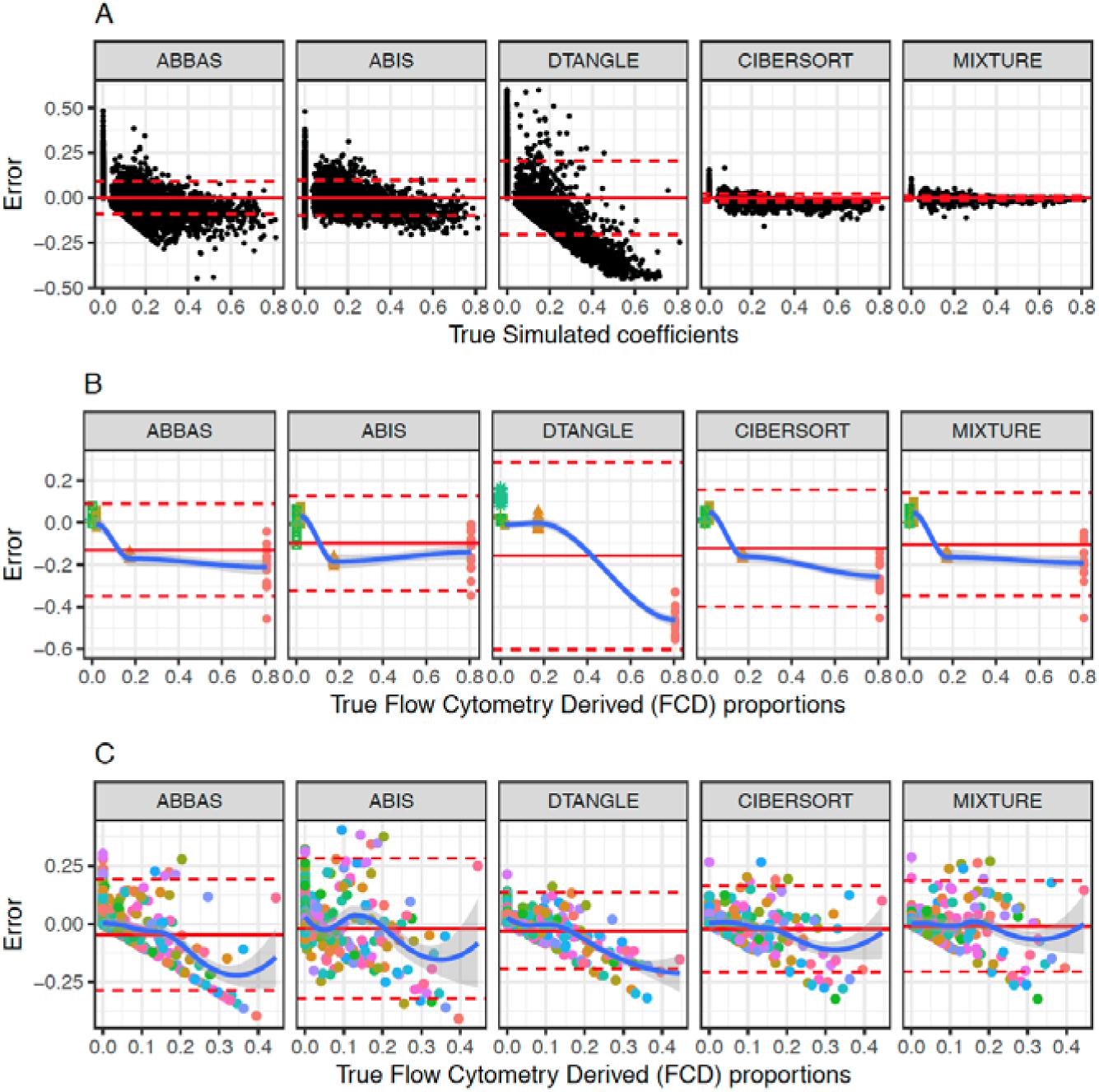
Bland-Altman plots to evaluate error bias between estimated and true (simulated and flow cytometry derived) proportion values. Horizontal continuous and dashed lines represent the overall mean and standard deviation errors. The light blue continuous line represents the smooth (loess) line to show prediction dependent bias. A: Estimation errors between predicted and simulated proportions, B and C: Estimation errors between predicted and flow cytometry derived cell-type proportions for FL and PBMC data respectively

### Evaluation over real mixture samples with flow cytometry derived proportions

Panels B and C from Figure 2 show the Bland-Altman plot of the differences between the estimated and B) flow cytometry derived (FCD) cell-types for the Follicular Lymphoma (FL) data, and C) FCD cell-types from Peripheral Blood Mononuclear Cells (PBMC) data (Scenario S2.1 and S2.2 respectively). For the first case (FL), DTANGLE tends to overestimate for low FCD values and underestimate towards higher proportion values, while for ABIS and MIXTURE methods resulted to be the most robust to estimation bias (no statistically significant difference between both methods), in accordance with results from Scenario S1.3. However, for the PBMC case, MIXTURE resulted to be the most robust against the overall mean difference between estimated and true flow cytometry derived proportions as well as shows to be more robust to bias towards high proportion values. It can be observed that the smooth curve (light blue continuous line), resulted closer to the error=0 value along the FCD axis for MIXTURE compared to the rest. In tables 1 and 2, the mean and standard deviation of the difference between estimated and flow cytometry derived cell-type proportions are shown (horizontal lines continuous and dashed lines, respectively, in Figure 2). It can be observed that ABIS and MIXTURE present the lesser overall bias (no significant difference between them, Wilcoxon paired test) for the FL data. However, MIXTURE resulted in being the one providing the lesser mean in PBCMC data sets for both FCD > 0 and FCD = 0. These differences were significantly smaller (paired Wilcoxon test p < 0.01) against the other methods.

**Table 1:** Summary statistics of the difference between estimated and Flow cytometry derived proportions for FL data

|  | FCD > 0 |  | FCD = 0 |  |
| --- | --- | --- | --- | --- |
|  | Mean | SD | Mean | SD |
| ABBAS | -0.129 | 0.123 | 0.043 | 0.076 |
| ABIS | **-0.098<sup>†</sup>** | **0.112** | 0.033 | 0.073 |
| DTANGLE | -0.157 | 0.222 | 0.052 | **0.039** |
| CIBERSORT | -0.122 | 0.144 | 0.040 | 0.056 |
| MIXTURE | -0.103<sup>*</sup> | 0.132 | 0.034<sup>*</sup> | 0.062 |

FCD: Flow cytometry-derived immune cell proportions. \* Statistically significant against the other methods except for ABIS (Wilcoxon paired test, $p < 0.001$ ). †: no significant differences against MIXTURE

**Table 2:**
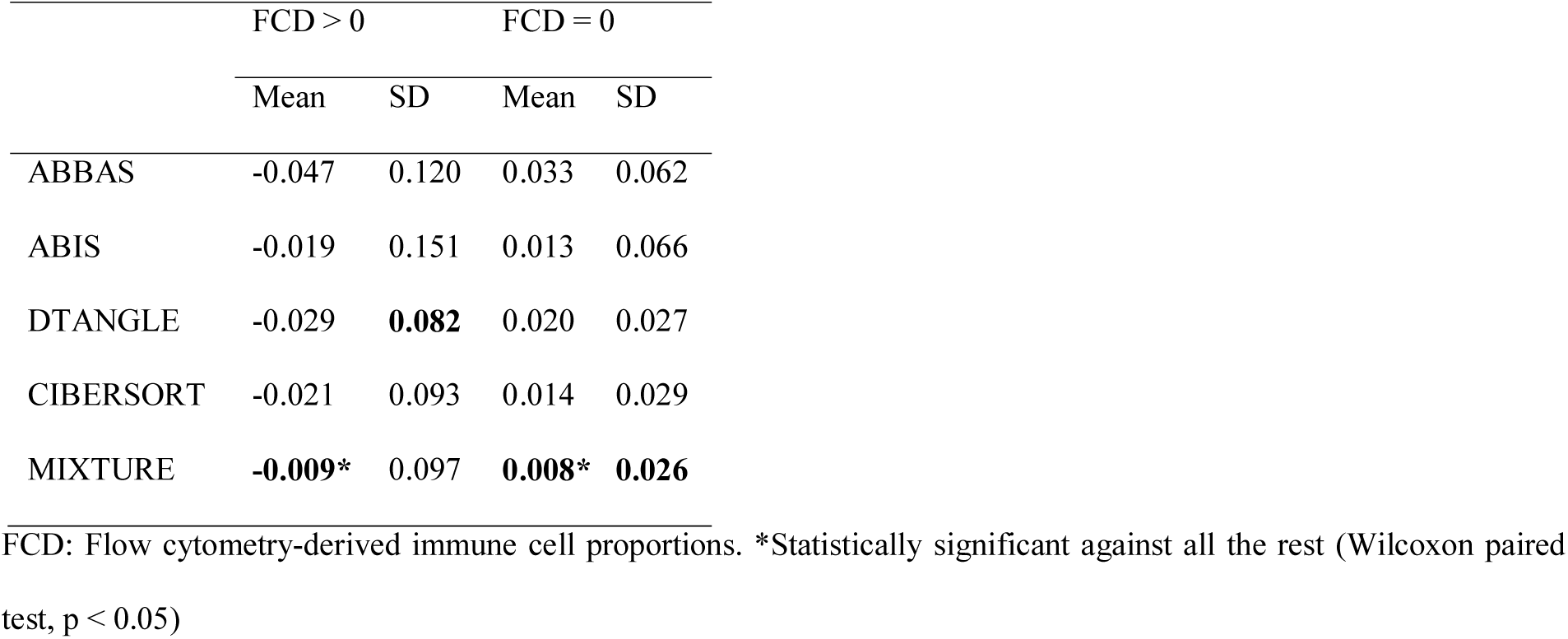
Summary statistics of the difference between estimated and Flow cytometry derived proportions for PBMC data

|  | FCD > 0 |  | FCD = 0 |  |
| --- | --- | --- | --- | --- |
|  | Mean | SD | Mean | SD |
| ABBAS | -0.047 | 0.120 | 0.033 | 0.062 |
| ABIS | -0.019 | 0.151 | 0.013 | 0.066 |
| DTANGLE | -0.029 | **0.082** | 0.020 | 0.027 |
| CIBERSORT | -0.021 | 0.093 | 0.014 | 0.029 |
| MIXTURE | **-0.009\*** | 0.097 | **0.008\*** | **0.026** |

FCD: Flow cytometry-derived immune cell proportions. \*Statistically significant against all the rest (Wilcoxon paired test, $p < 0.05$ )

### Applying MIXTURE on Breast Cancer TCGA data: Revealing the role of tumor-associated macrophages (TAMs)

Tumor-associated macrophages (TAMs) are abundant inflammatory cells in the tumor microenvironment capable of orchestrating different stages of breast cancer development. TAMs participate in the tumor angiogenesis, matrix remodeling, invasion, immunosuppression, metastasis, and chemo-resistance in breast cancer (Velaei et al. 2016). Several clinical studies indicate an association between the high influx of TAMs in tumor and the poor prognosis of hepatocellular, ovarian, cervical, breast cancer, among others (Qiu et al. 2018). Moreover, it has been revealed that M2 macrophages play a significant role in tumor development. This advocates for the efficient identification of the ITME to discover potential therapeutic targets that may help in the development of new therapies targeting tumor-associated macrophages (Hammerl et al. 2018).

Since all methods except those based on v-SVR present collinearity problems as well as high overall standard deviation errors, we applied CIBERSORT and MIXTURE to 1095 primary tumor (including 113 normal tissue) BRCA TCGA subjects (Scenario S3.1) to explore the ITME and its association with the outcome and PAM50 intrinsic subtypes. The subjects were confidently classified into the five PAM50 intrinsic breast cancer subtypes employing the PBCMC algorithm (18% Basal-like, 11% Her2-Enriched, 12% Luminal B, 25% Luminal A, 25% Not Assigned and 9% Normal-Like). We found that MIXTURE provides between 0 and 11 cell-types while CIBERSORT between 8 and 17, suggesting cell-type overestimation of the latter one. In Table 3, the proportion of samples for which each cell-type was identified is shown for the tumors samples. It is possible to observe that for most of the cell types MIXTURE identifies much fewer samples. However, MIXTURE identifies cell types known to be highly related to breast cancer survival with better discrimination power than CIBERSORT (see Figure 3), such as M2-macrophages, present in 97.72% of the tumor samples, follicular T-helper cells (93.79%), M0-macrophages (79.00%) and M1-macrophages (77.63%). They were the cell types more frequently found in breast cancer samples (see Table 3). It was described that tumor-associated macrophages (TAM) exhibit high plasticity in response to various external signals and play critical roles in innate and adaptive immune responses controlling numerous ITME factors strongly associated with clinical outcome (Choi et al. 2018).

**Table 3:**
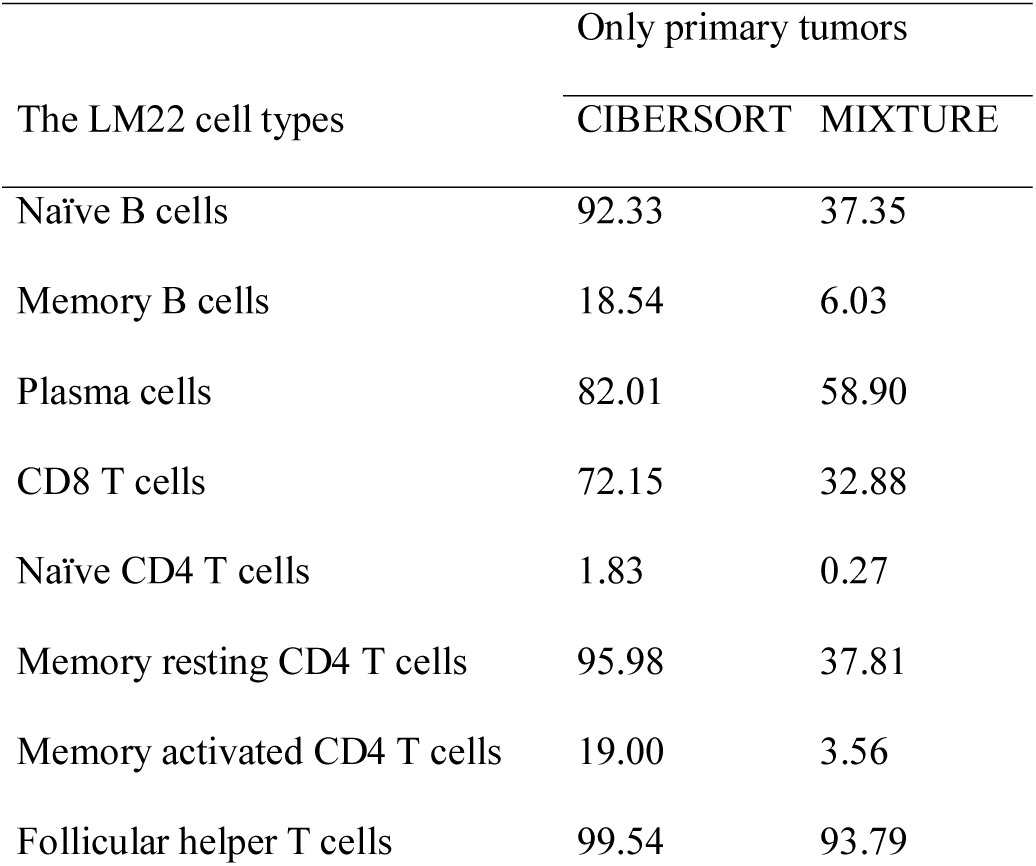
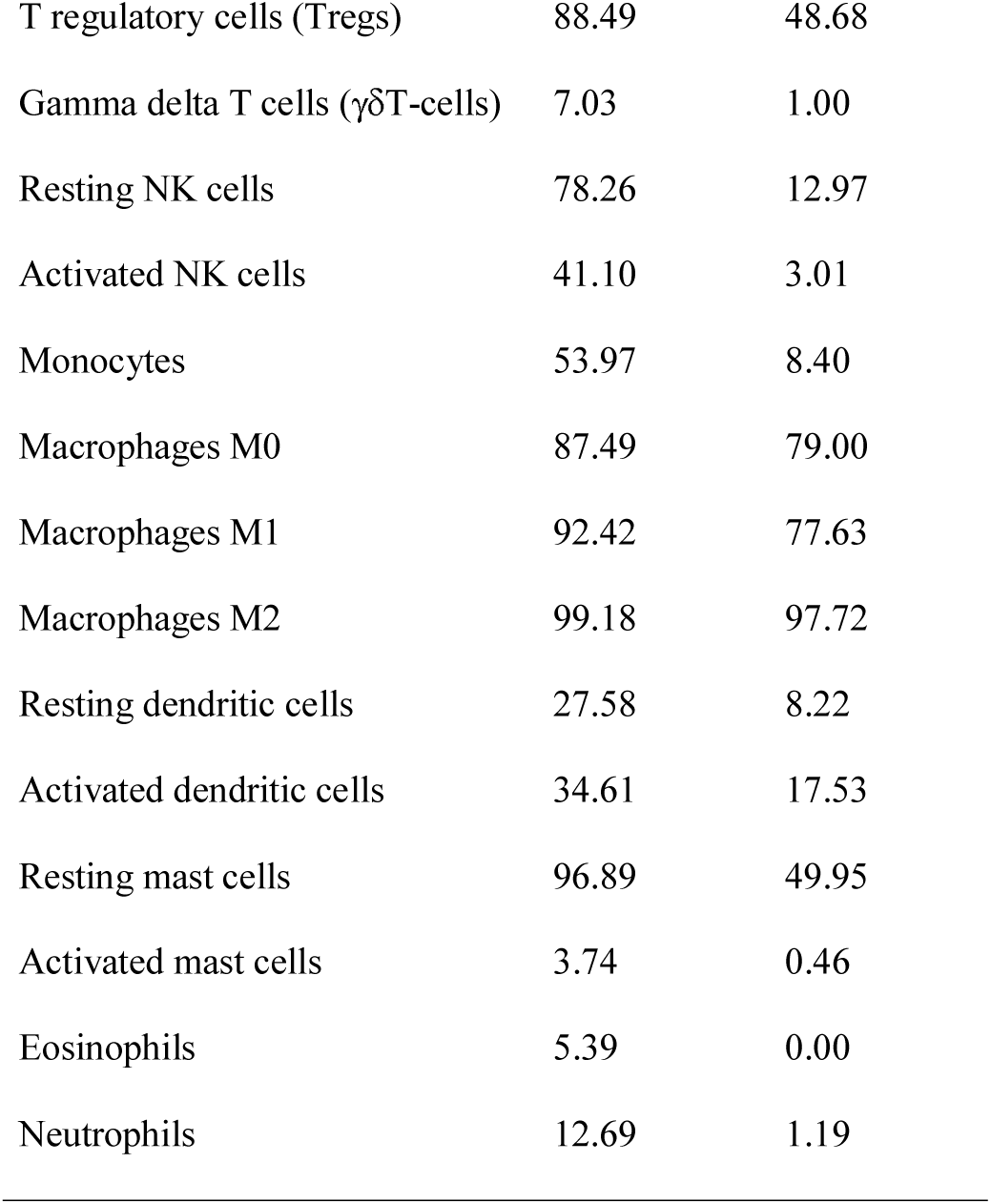
Percentage of cell types identified in the primary tumor TCGA BRCA cohort by CIBERSORT and MIXTURE, respectively.

| The LM22 cell types | Only primary tumors |  |
| --- | --- | --- |
|  | CIBERSORT | MIXTURE |
| Naïve B cells | 92.33 | 37.35 |
| Memory B cells | 18.54 | 6.03 |
| Plasma cells | 82.01 | 58.90 |
| CD8 T cells | 72.15 | 32.88 |
| Naïve CD4 T cells | 1.83 | 0.27 |
| Memory resting CD4 T cells | 95.98 | 37.81 |
| Memory activated CD4 T cells | 19.00 | 3.56 |
| Follicular helper T cells | 99.54 | 93.79 |
| T regulatory cells (Tregs) | 88.49 | 48.68 |
| Gamma delta T cells ($\gamma\delta$T-cells) | 7.03 | 1.00 |
| Resting NK cells | 78.26 | 12.97 |
| Activated NK cells | 41.10 | 3.01 |
| Monocytes | 53.97 | 8.40 |
| Macrophages M0 | 87.49 | 79.00 |
| Macrophages M1 | 92.42 | 77.63 |
| Macrophages M2 | 99.18 | 97.72 |
| Resting dendritic cells | 27.58 | 8.22 |
| Activated dendritic cells | 34.61 | 17.53 |
| Resting mast cells | 96.89 | 49.95 |
| Activated mast cells | 3.74 | 0.46 |
| Eosinophils | 5.39 | 0.00 |
| Neutrophils | 12.69 | 1.19 |

**Figure 3.**
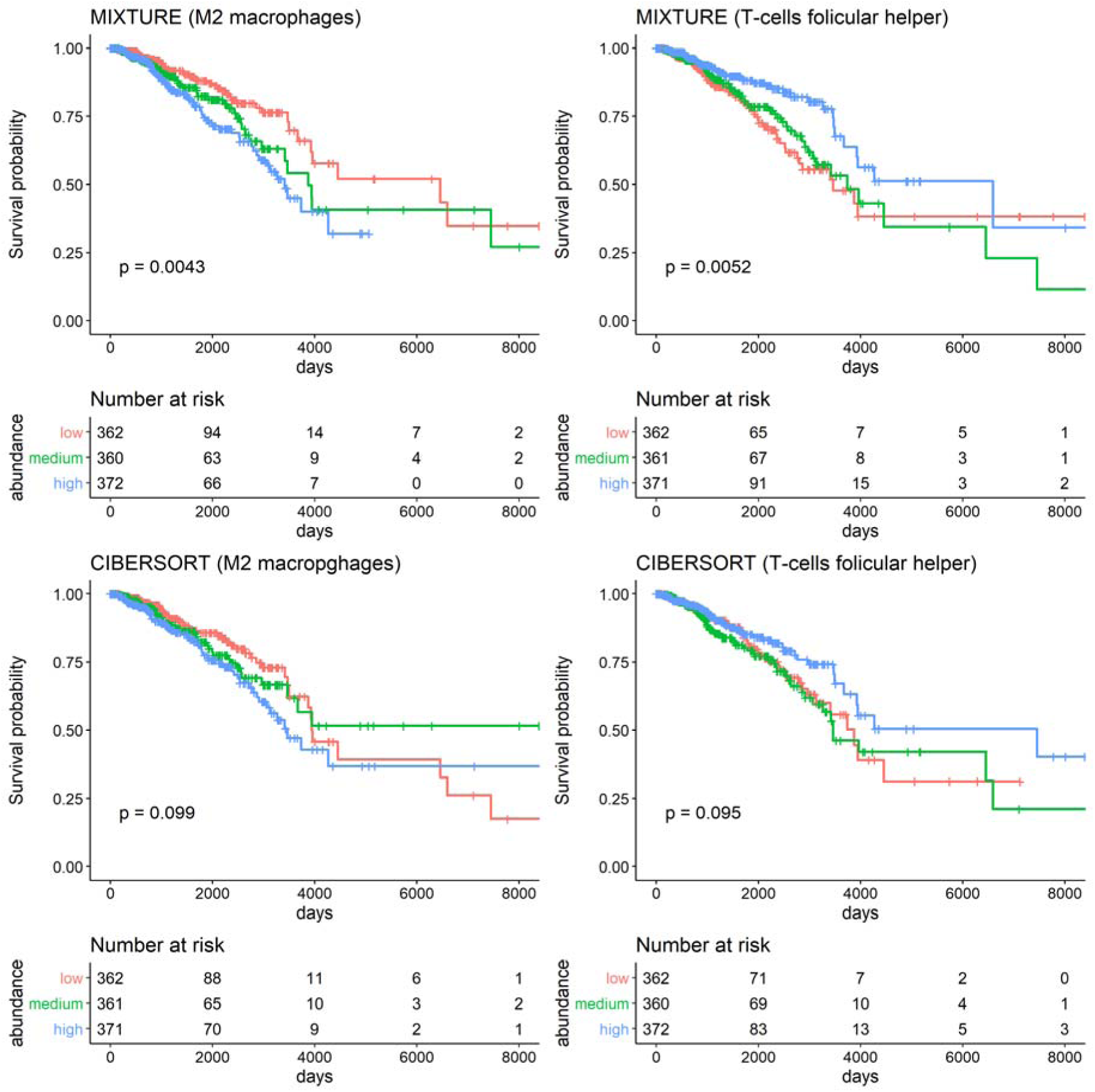
Kaplan Meier plots representing patient survival and immune infiltrate M2-macrophages (left panels) and follicular T-Helper cells (right panels) for the BRCA TCGA data by both methods (MIXTURE, top panels and CIBERSORT bottom panels). Each cell type proportion was split into three quantile levels (Low ≤ quantile 0.33, quantile 0.33 < Middle ≤ quantile 0.66, and High ≥ quantile 0.66). n=1095 (primary tumor samples)

By splitting estimated proportions into three percentile levels (Low=[0,0.33), Middle=[0.33,0.66) and High=[0.66,1]}) we found that M2-macrophages and follicular T-helper cells (FTHC) cell types resulted associated to worse survival independently of PAM50 only by MIXTURE estimation (p<0.01 for both cell types). These findings are supported by immunohistochemical staining of M2 and FTHC markers previously described by (Tiainen et al. 2015) and (Gu-Trantien et al. 2013) suggesting the appropriateness of MIXTURE estimations. For the Basal subtype, M2-macrophages were also strongly associated with outcome, but MIXTURE tends to stratify better the population reaching a smaller significance *p*-value (0.044 and 0.032 for CIBERSORT and MIXTURE respectively). When analyzing subjects by ER status, MIXTURE was the only method able to identify the association with survival for ER+ but only comparing Low vs High M2-Macrophages levels (p=0.036). The Low, Middle and High percentile M2-Macrophages levels were associated with survival for both methods in ER-case (P=0.016 and p=0.009 for MIXTURE and CIBERSORT respectively). High levels of M1-macrophages were found to be associated with a better prognosis for ER-subjects (p=0.008) meanwhile Low vs. High levels of FTHC were associated with worse prognosis for these subjects (p=0.038) only by MIXTURE. The results are in agreement with what was previously reported by Ali and colleagues (Ali et al. 2016) for a more extensive breast cancer population. It has been previously reported that targeting the reprogramming programs leading from M2 toward M1 macrophages phenotypes would be an efficient way to promote tumor regression which can be achieved through therapies including chemotherapy, immunotherapy, and radiotherapy (Genard et al. 2017), suggesting the importance of accurate TAM estimates. MIXTURE results regarding the behavior of M2, M1 macrophages and FTHC are in agreement with those previously found for the different breast cancer subtypes (Gu-Trantien et al. 2013; Gu-Trantien and Willard-Gallo 2017). Thus, MIXTURE represents a reliable and reproducible platform to estimate the nature and magnitude of particular immune cell populations in the tumor microenvironment.

### MIXTURE analysis reveals a differential immune infiltrate on HNSCC that correlates with PFS and OS

Head and Neck tumor development is closely related to the host immune system where the use of immunotherapy in the management of metastatic HNSCC may pave the way for future treatment due to their promising results (Schoenfeld 2015). These results advocate the study of the ITME and outcome association. For instance, Rickman *et al*, have predicted future metastasis development and characterized molecularly head and neck squamous cell carcinoma (HNSCC) based on transcriptome and genome analysis data (Rickman et al. 2008). The authors investigated the propensity for subsequent distant metastasis in head and neck squamous cell carcinoma HNSCC using 186 primary tumors from patients initially treated by surgery that developed (M) or did not develop (NM) metastases as the first recurrent event. Although the authors found that the most significantly altered transcripts in M versus NM were associated with metastasis-related functions, including adhesion, motility and cell survival, the study did not address the role of the immune infiltrate in the development of future metastasis. Here, we have inferred the immune infiltrate by CIBERSORT and by MIXTURE analysis of transcriptomic data of 41 biopsies from HNSCC patients who metastasized (M) vs. 40 biopsies from HNSCC patients who did not present metastasis (NM). We found that M2-macrophages were significantly increased in NM patients by CIBERSORT (p=0,034) while MIXTURE did not find a statistical significance between the M and NM cohorts (Figure 4A). The protumorigenic role of M2-macrophages has been widely reported and the literature does not support the increased proportion of this cell population in NM patients. Oppositely, while CIBERSORT did not find a differential infiltrate related to the monocyte infiltrate in the M vs. NM cohorts, MIXTURE found that the monocyte population was significantly increased in the metastatic (M) group (Figure 4B). This finding is supported by a previous study in HNSCC showing that patients with high monocyte frequency had lower survival with 8% 5-year overall survival (OS) compared to 65% 5-year OS for patients with low activation levels (Aarstad et al. 2015). Although this study was performed on peripheral blood, it shows a link between increased monocyte activation and decreased survival in HNSCC patients.

**Figure 4.**
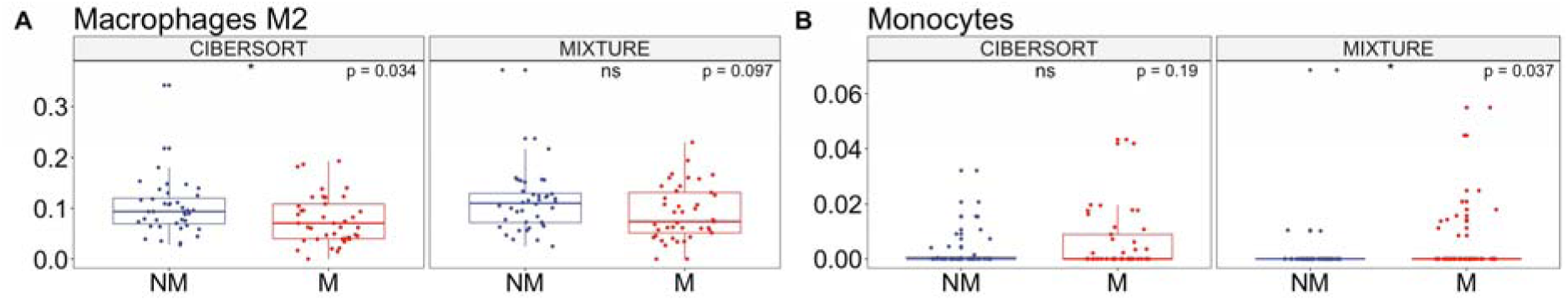
Box plots representing relative cell abundance of M2-macrophages and monocytes in biopsies of HNSCC patients who metastasized (M) vs. patients who did not present metastasis (NM) calculated by CIBERSORT or MIXTURE. M, n=41, NM, n=40.

### Applying MIXTURE to infer the immune infiltrate of melanoma biopsies and response to immunotherapy

Immunotherapy based on immune checkpoint blockade can induce durable clinical responses in melanoma patients. However, objective clinical responses are only observed in a small proportion of patients. Given the high financial costs and potential toxicities associated with these therapies, there is an urgent need to identify new, reliable biomarkers to distinguish better cancer patients likely to respond to immunotherapy. Riaz and colleagues (Riaz et al. 2017) assessed genomic changes in tumors from 68 patients with advanced melanoma, who progressed on the anti-CTLA-4 drug ipilimumab or were ipilimumab naive, before and after they commenced anti-PD-1 therapy with nivolumab. We analyzed public transcriptomic data from this cohort (Riaz et al. 2017) to study the tumor immune infiltrate before and during immunotherapy treatment in patients responding (Responders) and not-responding (Non-responders) to anti-PD-1 therapy using the CIBERSORT and MIXTURE algorithms.

Figure 5 shows that both methods revealed changes in distinct immune cell subsets before and during nivolumab therapy in responders and non-responders. However, only MIXTURE analysis revealed that M2-macrophage infiltrate decreased in biopsies from responding patients while CIBERSORT analysis only showed a trend that did not reach statistical significance (Figure 5A). This difference could be explained by the CIBERSORT limitation of underestimating highly represented immune cell populations. Vast evidence indicates that M2-macrophages are key players components of the immuno-suppressive tumor microenvironment (Cassetta and Kitamura 2018; DeNardo and Ruffell 2019) and several M2-macrophage targeting strategies (depletion, reprogramming, and targeting functional molecules) have been proposed to enhance the efficacy of the immune checkpoint inhibition. Tumor-associated macrophages were suggested as important therapeutic targets to enhance the efficacy of checkpoint blockade immunotherapies (Mantovani et al. 2017) thus its accurate identification turns crucial in understanding their tumor-associated activities and the design of novel anticancer agents. Also, the M2-macrophage population decrease in biopsies from patients responding to immunotherapy as previously shown for anti-CTLA-4 therapy (Yu et al. 2016).

**Figure 5.**
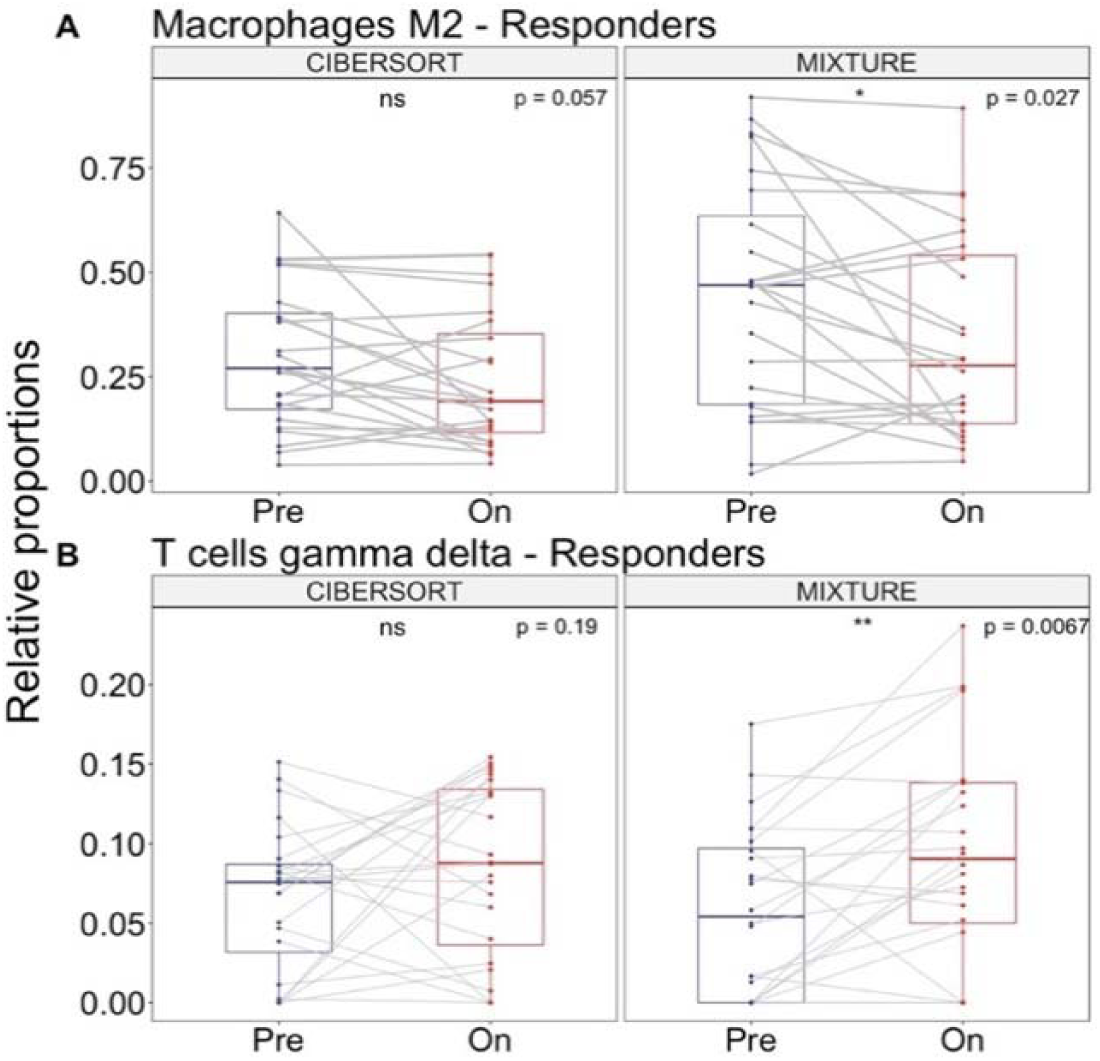
Paired analysis of relative cell abundance of M2-macrophages and γδT-cells in pre-treatment vs. on-treatment melanoma biopsies calculated by CIBERSORT or MIXTURE from patients responding to anti-PD-1 immunotherapy; n=24.

Interestingly, only MIXTURE analysis revealed an increase in the proportion of γδT-cells in biopsies of responding patients (Figure 5B) while CIBERSORT analysis did not reach statistical significance. The relevance of this finding is supported by previous literature showing the cytotoxic potential of γδT-cells against tumor cells (Silva-Santos et al. 2015; Wu et al. 2017; Zou et al. 2017; Lo Presti et al. 2018; Zhao et al. 2018). Biologically, vast publications support the antitumor role of γδT-cells by direct or indirect ways. γδT-cells recognize antigens shared by tumor cells in an MHC-II independent manner and mediate antitumor effects by the perforin-granzyme pathway, through the ligands TRAIL and FasL (death-inducing ligands), via antibody-dependent cellular cytotoxicity and production of IFN-γ and TNF-α with cytotoxic activity against tumor cells. This effect may occur directly or indirectly via stimulating other immune cell types such as macrophages or dendritic cells (Lo Presti et al. 2018; Zhao et al. 2018). Moreover, the intratumoral infiltrate of γδT-cells has been identified as the most significant predictor of survival in some cancer types including ovarian cancer, in which these cells display critical immunosuppressive effects by releasing high amounts of pro-tumorigenic galectin-1 (Rutkowski et al, 2015). In the last years, these cells have become critical cells with both effector and immunosuppressive activity in cancer settings. These findings suggest that MIXTURE analyses are more robust and meaningful.

## Discussion

In this study, we present a new digital cytometry method, MIXTURE, for accurate estimation of cell-type content of bulk tissues which provide a meaningful understanding of the ITME behavior in cancer samples. The MIXTURE algorithm implements a novel approach to handle the overestimation of the amount of present ITME cell types utilizing a noise constraint threshold to prevent floating point errors jointly with the Recursive Feature Selection (RFE) process which improves the known robustness of the v-SVR to collinearity. In this way, it provides a robust estimation of the tumor immune microenvironment content, defined from a given and validated molecular signature matrix. The added advantage of providing accurate estimates of the cell-types composition positively impacts the consequent improvement on the accuracy of estimated cell types proportions. The MIXTURE algorithm also diminished the estimation bias towards high proportion values, shown by the other evaluated methods. This estimation bias may negatively impact the estimation process hiding relevant biological information. The appropriateness of MIXTURE was verified using simulated and benchmark flow cytometry derived content data. Also, by using real cancer data, MIXTURE reveals that M2-macrophages proportions levels correlate with poor prognosis in BRCA TCGA samples independently of the subtype and also for their Basal and ER+ subtypes as well as their role in the immunotherapy response in melanoma samples. These associations against survival and therapy responses were previously identified in such cancer types by other authors and techniques. However, they were not found by using CIBERSORT proportion estimates.

The same happened with follicular T-Helper cells and γδT cells, where MIXTURE estimates allow to identify known and well-described associations with survival for BRCA and with therapy response for melanoma. On the contrary, CIBERSORT found an association of their M2-macrophages estimated levels with the development of metastasis in HNSCC meanwhile MIXTURE not. However, we were not able to find supporting bibliography evidence of such association. Conversely, MIXTURE provides monocyte levels and their association to metastasis development, which has been previously reported. All these differences may be due to both floating-point errors and an incomplete feature selection process as well as collinearity that may lead to missing estimation of cell types number and proportion levels. Collinearity problems have been recognized strongly affecting the estimation of cell type contents in digital cytometry and has been addressed in tools like TIMER (which use OLS), but the method should be set for each molecular signature trough a specific correlation analysis thus it cannot be easily extended to other molecular signature matrices. In this context, SVR method has been proven to be robust to collinearity in several scenarios (Fernandez et al. 2011; Newman et al. 2015; Newman et al. 2019). However, the noise constrained RFE implemented in MIXTURE improves the collinearity robustness of the v-SVR implemented in CIBERSORT, with the consequent advantage of diminishing the amount of false cell type detections and estimation bias. The MIXTURE algorithm was developed both as an R code and as a shiny application available for downloading from GitHub repository (a guide for installation, use, and application of MIXTURE to other data sets is available in Supplementary Material). It can be easily implemented as a previous digital cytometry step for the determination of cell-type expression profiles from bulk tissues as used by CIBERSORTx (Newman et al. 2019).

## Methods

### Mixture: a novel algorithm to infer the immune infiltrate in tissue samples

The linear deconvolution of the cell types present in a gene signature matrix (X), holding *N* genes for *k* cell types, associated with the components of a mixture of cell types present in a tumor gene expression profile (***Y***), involves solving the following regression model equation **Y= X ⋅ B^T^**, where the proportions for all cell types in the mixture sample is represented by the column tor **B** = {β_*j*_≥ 0 ^ ∑ β_*j*_ = 1 ∀*j* = 1,…,*k*}, i.e. the vector of regression coefficients satisfying both, the non-negativity and sum-to-one constraints (Avila Cobos et al. 2018). Among the several algorithms developed for estimating **B**, the most competing ones (Li et al. 2016; Hunt et al. 2018) are the ordinary least squares (OLS), initially presented by Abbas and colleagues (Abbas et al. 2009) [ABBAS method currently used in TIMER tool by (Li et al. 2016)], the ν-support vector regression (ν −SVR) implemented in CIBERSORT (Newman et al. 2015), the recently presented algorithms: DTANGLE (Hunt et al. 2018) based on multivariate logistic function, and the ABIS method based on Robust Linear Model [rlm, (Monaco et al. 2019)].

The ABBAS, ABIS and CIBERSORT methods, estimate the regression coefficients utilizing OLS, rlm or v-SVR, the later a machine learning algorithm claimed to be robust to multicollinearity, yielding 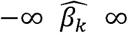. To satisfy β_*k*_≥ 0 ∀ *k*, in (Abbas et al. 2009), the columns of ***X*** associated with those 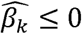 are iteratively removed until all the estimated regression coefficients resulted positive (A recursive feature selection approach). The CIBERSORT method provides a set of 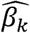 coefficients obtained using an optimized v-SVR by selecting *ν* (i.e., a regularization parameter controlling the number of support vectors included in the solution) over three fixed values (ν = 0.25,0.5 and 0.75). It chose the one that minimizes the root mean squared error (RMSE) after setting as null those 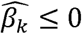. Finally, both algorithms normalize the resulting vector of regression coefficients as 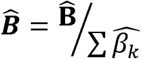 to satisfy ∑β_*k*_ = 1. On the other hand, the ABIS method directly estimates 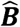 utilizing the Robust Linear Model (rlm) without setting as null those 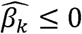 nor normalizing 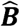 thus not forcing to satisfying the non-negativity and sum-to-one constraints (Avila Cobos et al. 2018).

These methods work with non-log RNA-Seq data or one-color microarray technology, as proposed by (Zhong and Liu 2011), for the preservation of the underlying linear relationship between the observed gene expression profile and the gene signature matrix. On the contrary, DTANGLE, provides proportion estimates by mapping the weighted average difference between the signature and the sample mixture profiles, in the log2 scale, into the unit interval [0,1] by a multivariate logistic function (Hunt et al. 2018).

Our newly developed algorithm, MIXTURE, re-estimates **B** by iteratively removing the columns of ***X*** associated with null coefficients from the regression step, after all estimated coefficients resulted not-null, as in ABBAS, but using the v-SVR approach as a regression function similar to CIBERSORT. However, we observed that floating-point errors affect the inequality comparison 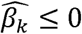 used to define the null coefficients, which produce overestimation of the number of positive coefficients, thus negatively affecting the feature selection step. To avoid these floating-point errors, MIXTURE defines a noise constraint threshold ∆ (equal to 0.007 and chosen through a simulation process (see supplementary material). In this way, active coefficients (i.e. not-null ones) are recognized as those normalized β_k_ > ∆, thus identifying the cell types present in the sample (see Supplementary Table 1 for MIXTURE algorithm implementation details).

The ABBAS least square method was implemented following the source code of (Li et al. 2016). The CIBERSORT method was run from the web site https://cibersort.stanford.edu in February 2019, the ABIS method was implemented using the code provided at https://github.com/giannimonaco/ABIS and for DTANGLE the code implemented by (Hunt et al. 2018) was run from their “DTANGLE” R library.

### Statistical Analysis

The Wilcoxon test was used to compare the estimated coefficient between methods (paired or unpaired accordingly) with p < 0.05 as the significance threshold value. The Bland-Altman Statistical method for assessing agreement between clinical measurements (Bland and Altman 1986) was used to compare estimated proportions against the true simulated or flow cytometry derived ones (Fernandez et al. 2001). The mean ± standard deviation was used to show the overall bias when appropriate.

### Data Access

All data access description is detailed in supplementary material and available at https://github.com/elmerfer/MIXTURE.App

## Acknowledgments

Work in our laboratories is supported by grants from Agencia Nacional de Promoción Científica y Tecnológica (PICT 2016-2130 to MRG), Harry J. Lloyd Foundation Trust to MRG, The Argentinean National Council of Scientific Research (CONICET) to EAF, the Universidad Católica de Córdoba to EAF and the National Cancer Institute (Argentina) to YDM and MRG.

